# GSK3 inhibition improves skeletal muscle function and whole-body metabolism in the severe DBA/2J *mdx* mouse model

**DOI:** 10.1101/2022.02.16.480726

**Authors:** Bianca M. Marcella, Briana L. Hockey, Jessica L. Braun, Kennedy C. Whitley, Mia S. Geromella, Ryan W. Baranowski, Colton J.F. Watson, Sebastian Silvera, Sophie I. Hamstra, Luc J. Wasilewicz, Robert W.E. Crozier, Amelie Marais, Rene Vandenboom, Brian D. Roy, Adam J. MacNeil, Rebecca E.K. MacPherson, Val A. Fajardo

## Abstract

Duchenne muscular dystrophy (DMD) is a severe X-linked muscle wasting disorder that affects 1 in 5,000 males worldwide^1^. It is caused by the absence of functional dystrophin, which compromises muscle integrity, leading to progressive muscle wasting and weakness^2^. Glucocorticoids are the standard of care for patients with DMD as they delay the loss of ambulation by an average of 3 years^3^; however, they are also associated with adverse effects such as insulin resistance and increased risk of type 2 diabetes^4^. Thus, alternative therapeutic options should be explored. Here, we show that treating the DBA/2J *mdx* mouse with the glycogen synthase kinase 3 (GSK3) inhibitor, tideglusib, improved skeletal muscle function and insulin sensitivity, while also attenuating the hypermetabolic phenotype previously observed in these mice^5^. Furthermore, treating *mdx* mice with the GSK3 inhibitor, lithium, augmented the benefits of voluntary wheel running on insulin sensitivity and skeletal muscle function despite running half of the total distance compared to control-treated *mdx* mice. This is important given that some patients with DMD may not be able to engage in adequate amounts of physical activity. Thus, GSK3 inhibition alone or in combination with exercise can enhance skeletal muscle function and insulin sensitivity in *mdx* mice.

---

GSK3 is a serine/threonine kinase that was first identified for its role in regulating glycogen synthase in muscle^6^. There are two GSK3 isoforms, GSK3α and GSK3β, with the latter being the most expressed and active isoform found within muscle^7^. In addition to inhibiting glycogen synthase, GSK3 can phosphorylate and inactivate the insulin receptor, insulin receptor substrate, and trafficking regulator of the glucose transporter type 4 (GLUT-4)^8, 9^. Thus, GSK3 acts on several pathways to limit insulin-mediated glucose disposal, and its inhibition could be beneficial for DMD patients - especially those undergoing glucocorticoid therapy as they are at increased risk of developing type 2 diabetes^10–12^.

In addition to improving insulin sensitivity, we and others have shown that inhibiting GSK3 can increase muscle mass, strength, regeneration, and the proportion of slow oxidative fibers in muscle, which altogether can help attenuate dystrophic pathology in DMD boys^13–17^. This is because DMD is a condition characterized by excessive muscle wasting and weakness that particularly affects fast-type muscles as they are more susceptible to dystrophic pathology^18, 19^. To our knowledge, there has been no study investigating whether inhibiting GSK3 can alleviate DMD pathology; however, previous studies have shown that GSK3 inhibition can alleviate other muscle wasting conditions such as myotonic dystrophy^20, 21^, limb girdle muscular dystrophy^22^, and muscle unloading^13, 15^. Furthermore, GSK3 has been shown to be overactive in muscles obtained from the preclinical *mdx* mouse model for DMD – making GSK3 a plausible therapeutic target^23^. To test this further, we examined the effects of pharmacological GSK3 inhibition in the severe DBA/2J *mdx* mouse model for DMD^24–26^.

We first examined the effects of tideglusib treatment in young and old DBA/2J *mdx* mice. Tideglusib is a clinically advanced GSK3 inhibitor currently being tested in clinical trials for conditions such as supranuclear palsy, autism spectrum disorders, and myotonic dystrophy^27, 28^. In this study, we used a dose of 10 mg/kg body mass/day, which is in line with a 600 mg dose for a 60 kg patient. This dose has been used for patients with myotonic dystrophy and is proven to be highly tolerable with little adverse effects^20^. Mechanistically, the primary action of tideglusib is to irreversibly bind to GSK3, thereby locking it in an inactive state^29^. For the young cohort, tideglusib treatment started at 6-8 weeks of age for a total of 4 weeks, and as such, treatment occurred in the midst of the initial bout of myonecrosis found in DBA/2J *mdx* mice^25^. For the aged cohort, tideglusib treatment also lasted 4 weeks and was initiated at 26-28 weeks, which is an age characterized by a phase of heightened muscle regeneration^25^.

Muscle performance and fatiguability, measured as hangwire time normalized to body mass (e.g., impulse), was significantly lower in aged *mdx* mice compared with young *mdx* mice, irrespective of treatment (Fig. 1a). However, there was a main effect of tideglusib, indicating that treatment with the GSK3 inhibitor improved hangwire performance relative to vehicle treated mice (Fig. 1a). Similarly, twitch and tetanic specific force production in the extensor digitorum longus (EDL), a fast muscle that is weakened in the DBA/2J *mdx* mouse^24^, was significantly greater in young and old tideglusib treated *mdx* mice compared with vehicle treated *mdx* mice (Fig. 1b-c). We also found a significant main effect of tideglusib when assessing serum CK levels, indicating that tideglusib treatment lowered muscle degeneration (Fig. 1d). A significant main effect of age for serum CK (Fig. 1c) was also found, and this is in agreement with previous findings showing that aged DBA/2J *mdx* mice have lower levels of circulating CK when compared with young *mdx* mice^26^.

**Fig. 1.**
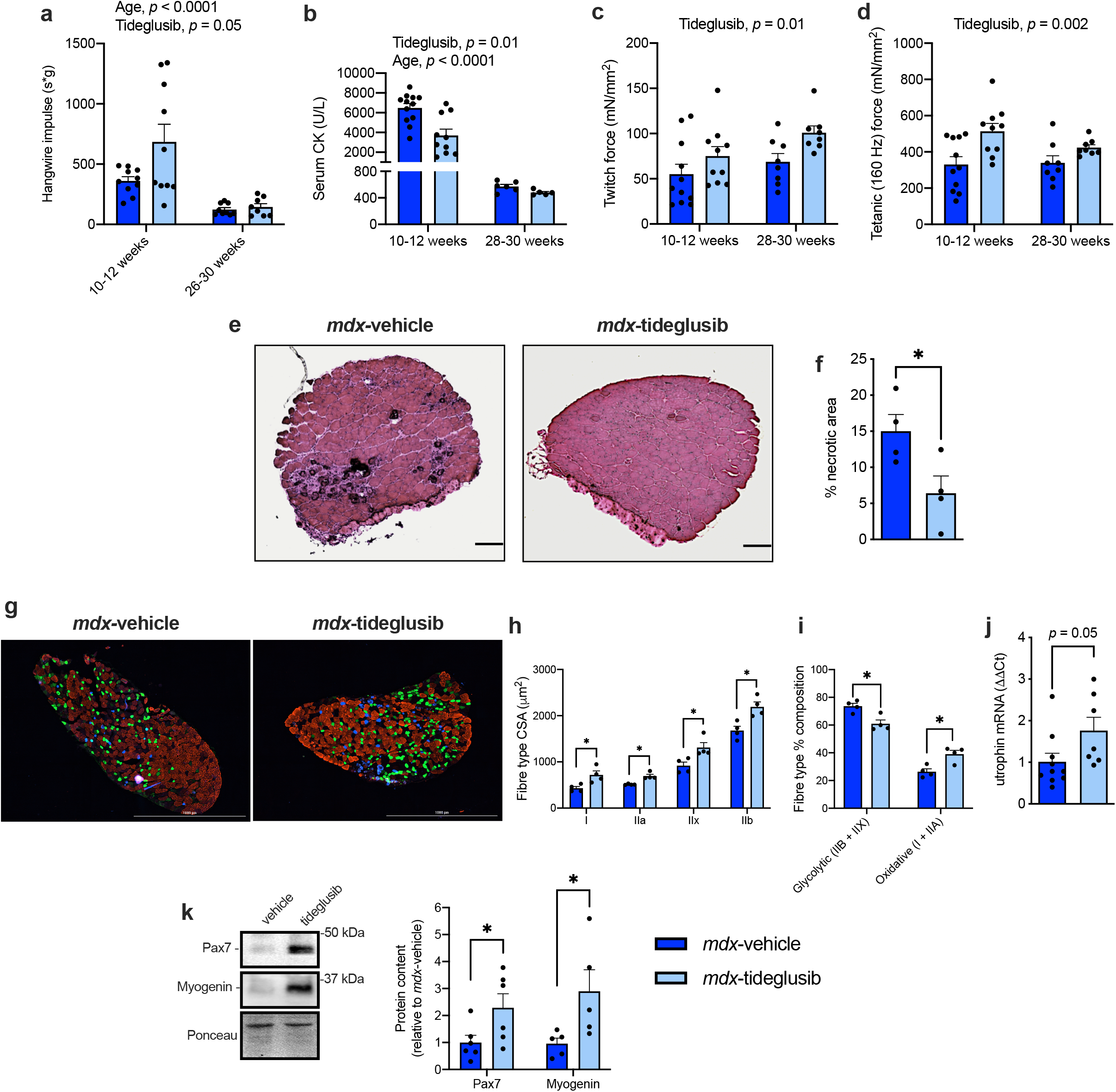
Tideglusib treatment improves muscle performance, reduces muscle necrosis, and promotes the oxidative fiber type in *mdx* mice. **a)** Hangwire impulse is lower in aged *mdx* mice and greater with tideglusib treatment in both young and aged *mdx* mice (n = 8-10 per group). **b-c)** Twitch and tetanic specific force production in the EDL is improved with tideglusib treatment (n = 8 – 11 per group). **d)** Serum creatine kinase (CK) levels are lowered with tideglusib treatment and with age (n = 5-12 per group). **e-f)** Muscle histological analysis with H&E staining shows a significant reduction in % necrosis in EDL muscles obtained from young tideglusib treated *mdx* mice (n = 4 per group). **g-i)** Tideglusib treatment increases myofiber cross-sectional area (CSA) and the proportion of oxidative fibers (type I and IIA) (n = 4 per group). Blue, type I fibers; green, type IIA fibers; red, type IIB fibers; unstained (black), type IIX fibers. **j**) Tideglusib treatment increases utrophin expression in *mdx* plantaris muscles (n = 7-10 per group).**k**) Tideglusib increases Pax7 and myogenin content in EDL muscles obtained from aged *mdx* mice. For **a-d**, a two-way ANOVA was used to assess the main effects of age and tideglusib treatment. Significant main effects are denoted in the text above. For **f,h-k**, a Student’s t-test was used. All values are mean ± SEM. \**p*<0.05.

We next examined the effects of tideglusib treatment on muscle histology in young *mdx* mice, given that this age is characterized by excessive muscle degeneration and necrosis^25^. Strikingly, we found that tideglusib treatment significantly lowered myonecrosis in the EDL (Fig. 1e-f). This result was accompanied by a significant increase in myofiber cross-sectional area (Fig. 1g-h), which is consistent with the known hypertrophic effect of GSK3 inhibition^16, 17^. Moreover, and in agreement with the repressive action of GSK3 on the oxidative fibre type^14, 17^, tideglusib treatment significantly lowered the proportion of glycolytic fibres (IIB and IIX) while increasing the proportion of oxidative (I and IIA fibres) (Fig. 1g and i). Further, plantaris muscles from tideglusib treated *mdx* mice had greater utrophin expression when compared with vehicle treated mice (Fig. 1j). Utrophin is a dystrophin homolog that can provide compensatory membrane stability in the absence of dystrophin, and it is found predominantly in oxidative muscle fibers^19^. This is because its expression is largely controlled by the calcineurin/nuclear factor of activated T-cell (NFAT) signaling axis^18, 19^.

Calcineurin is a Ca^2+^-dependent phosphatase that dephosphorylates NFAT leading to its nuclear localization where it can promote the expression of genes associated with the oxidative phenotype (including utrophin)^19^. In addition to regulating utrophin expression, calcineurin can also promote the expression of genes involved with muscle regeneration, and therefore, its activation provides multiple benefits that can attenuate dystrophic pathology in *mdx* mice^18, 30, 31^. Given that we and others have shown that GSK3 counteracts calcineurin signaling^14, 32, 33^, we attribute at least part of the benefits observed with GSK3 inhibition in *mdx* mice to the activation of calcineurin. With respect to muscle regeneration, we next assessed the protein levels of Pax7 and myogenin, which are markers of muscle regeneration^15^. We focused on aged mice for these measures since at this age, the *mdx* mice are known to be undergoing a period of heightened muscle regeneration^25^. Our results show that tideglusib treatment significantly increased the protein levels of Pax7 and myogenin in EDL muscles when compared with vehicle controls (Fig. 1k). These findings are strongly in line with previous work showing that GSK3 inhibition accelerates muscle regeneration^13, 15, 16^. Collectively, these data show that GSK3 inhibition in the DBA/2J *mdx* mouse can impart several benefits to muscle strength and quality by lowering muscle necrosis, increasing muscle size, and promoting the oxidative fiber type and muscle regeneration.

It has been previously established that *mdx* mice present with a hypermetabolic phenotype^5, 34^, expending a significant amount of energy on a daily basis to support the costs of both muscle degeneration and regeneration. In young and aged *mdx* mice, tideglusib treated groups had lower energy expenditure when compared with vehicle treated groups (Fig. 2a-b). This effect was not due to any changes in daily cage ambulation (Fig. 2c-d). Furthermore, the hypermetabolic phenotype in *mdx* mice has been previously shown to lower body fat content, resulting in less whole-body lipid supply, thereby raising the respiratory exchange ratio (RER)^34^. In our hands, DBA/2J *mdx* mice also had a higher RER when compared with WT mice (Extended Data Fig. 1). However, and perhaps cohesive with the reduction in energy expenditure afforded by tideglusib treatment, both young and aged tideglusib treated *mdx* mice had higher body fat content when compared with vehicle treated mice, though this effect was most prominent at 28-30 weeks of age (Fig. 2e-f). Moreover, tideglusib treated *mdx* mice had lower RER values when compared with vehicle treated mice (Fig. 2g-h).

**Fig. 2.**
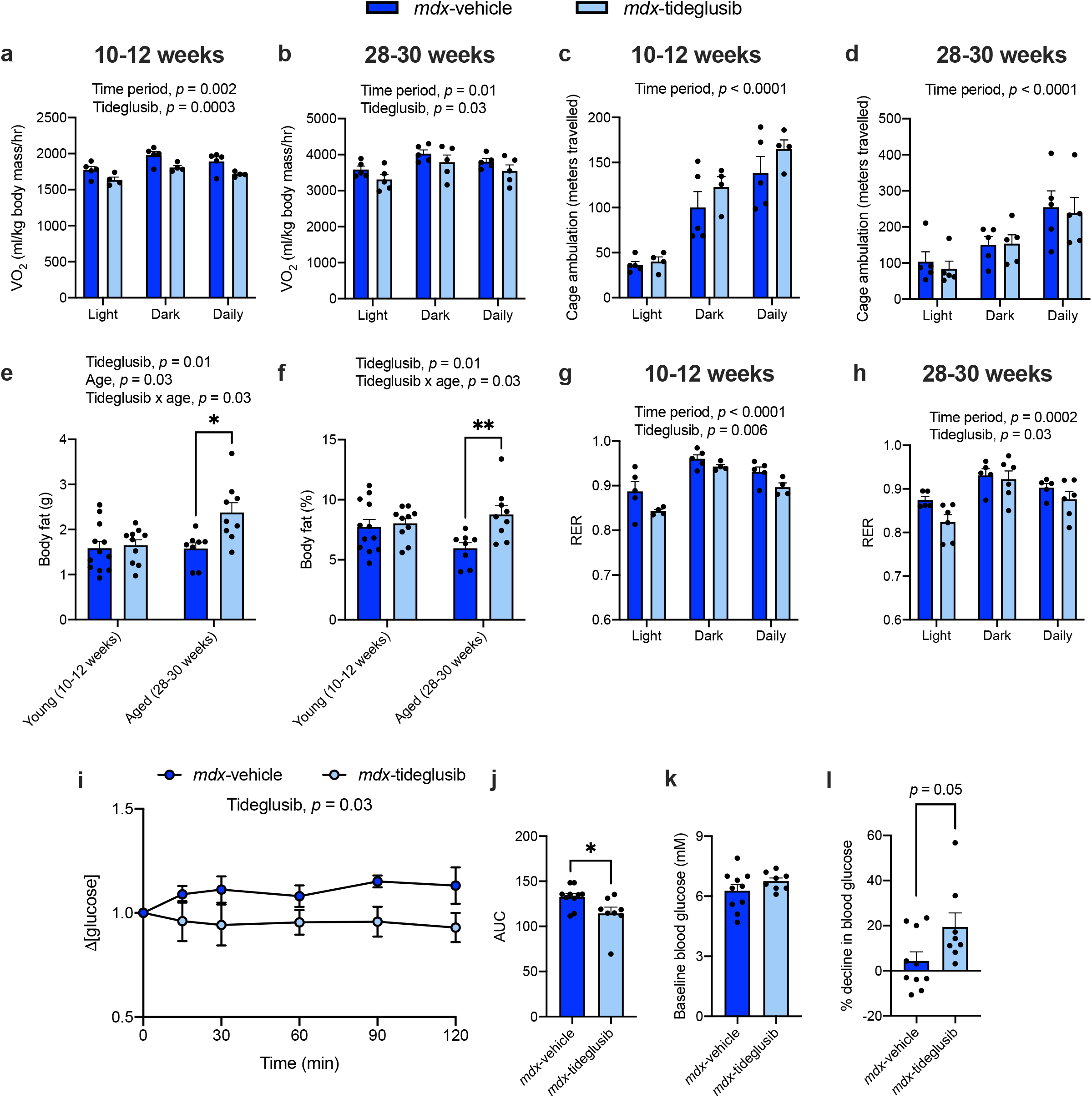
Tideglusib treatment attenuates several metabolic alterations typically found in *mdx* mice. **a-b)** Tideglusib lowers daily energy expenditure in young and aged *mdx* mice (n = 4-5 per group). **c-d)** Tideglusib treatment did not alter cage ambulation (n = 4-5 per group). **e-f)** Body fat content (g) and body fat composition (% of body mass) were elevated in aged *mdx* mice and in tideglusib treated young and aged *mdx* mice (n = 8-12 per group). **g-h)** Tideglusib lowers the respiratory exchange ratio (RER) in young and aged *mdx* mice (n = 4-5 per group). **i-l)** Insulin tolerance tests and corresponding area-under-the-curve (AUC) analysis, baseline glucose levels and % decline in blood glucose in 28-30 week old *mdx*-vehicle and *mdx*-tidgelusib mice (n = 8-10 per group). Glucose traces are displayed relative to baseline glucose levels. AUC were obtained from the normalized traces. There were no differences in baseline glucose levels. For **a-d**, a two-way ANOVA was used to assess the main effects of time period and tideglusib treatment. For **e-h**, a two-way ANOVA was used to assess the main effects of age and tideglusib treatment with a Tukey’s post-hoc in the event of a significant interaction. Significant main effects and interaction terms are denoted in the text above. For **j-l**, a Student’s t-test was used. All values are mean ± SEM. \**p*<0.05, \*\**p*<0.01.

In addition to the hypermetabolic phenotype, *mdx* mice are well-known to be insulin resistant^35^. Here, we show that vehicle treated *mdx* mice showed very little response to insulin during an insulin tolerance test, and many of these mice even displayed a slight increase in blood glucose levels after insulin injection (Fig. 2i-l). In contrast, tideglusib treated *mdx* mice were more responsive to insulin, leading to a ∼20% drop in blood glucose levels over time and a significant reduction in the area-under-the-curve (AUC, Fig. 2i-l). Altogether, our results show that tideglusib treatment improves muscle strength, lowers muscle necrosis, and enhances insulin sensitivity in DBA/2J *mdx* mice. Indeed, these results are likely related as skeletal muscle accounts for more than 90% of whole-body insulin-stimulated glucose removal^36^ and thus improving muscle health can positively impact glucose regulation.

Exercise is also a powerful stimulant for insulin sensitivity and overall muscle health, though its role in DMD disease management has been controversial^37^ and for the most part, the recommended exercises have been limited to mild or moderate intensity regimes^38, 39^. This highlights a potential opportunity to develop adjuvant therapies that can be combined with low-intensity forms of exercise. Thus, we next investigated the combined effects of exercise and GSK3 inhibition on muscle form and function in young *mdx* mice. Voluntary wheel running (VWR) was chosen as the mode of exercise given its low intensity and minimal stress added to the mouse^40^. To further minimize stress, we treated the mice with the GSK3 inhibitor, lithium (Li), as it could be provided easily through drinking water rather than performing oral daily gavage as with tideglusib. Li is a natural inhibitor of GSK3 and is most commonly used for the treatment of bipolar disorder^41^; however, its clinical use must occur within a narrow therapeutic range of 0.5-1.0 mM serum Li concentration to avoid potential adverse effects on various tissues^41^. Here, we treated *mdx* mice with a low dose of lithium chloride (50 mg/kg/day via drinking water) for 6 weeks, which resulted in a serum Li concentration of 0.15 ± 0.05 mM (n = 9).

We examined the effects of VWR with and without Li supplementation on EDL and soleus muscle contractility, given that the soleus muscle is responsive to VWR^42^. In the soleus, twitch and tetanic force were significantly depressed in *mdx-*SED mice when compared with WT mice (Fig. 3a-b). Though VWR appeared to improve soleus force production in *mdx* mice, it did not lead to any significant differences when compared with *mdx*-SED mice (*p* = 0.07 and 0.09 for twitch and tetanic force comparisons, respectively). Furthermore, the *mdx*-VWR soleus force outputs were still significantly lower when compared with WT mice (Fig. 3a-b). In contrast with this, the combination of VWR and Li provided additional benefits to soleus force production that were significantly different from *mdx*-SED mice and were no longer different from WT mice (Fig. 3a-b). In the EDL, twitch and tetanic force were significantly lower in all *mdx* groups compared with WT mice; however, force output was improved with Li supplementation such that twitch and tetanic force production were greater in the *mdx*-VWRLi group when compared with *mdx*-VWR (Fig. 3c-d). These improvements in soleus and EDL contractile function observed in the *mdx*-VWRLi mice was accompanied by reduced muscle necrosis compared with *mdx*-SED mice (Fig. 3e-f). This is consistent with our tideglusib experiments. Furthermore, a significant main effect of muscle type for muscle necrosis was found, indicating that across treatment groups, the EDL had greater % necrosis when compared with the soleus (Fig 3f). This was expected as glycolytic muscles are more susceptible to dystrophic pathology^18, 19^. We also found that VWR increased utrophin expression in the quadriceps, which is consistent with previous work^43^, however, Li supplementation provided no added benefit (Fig. 3g).

**Fig. 3.**
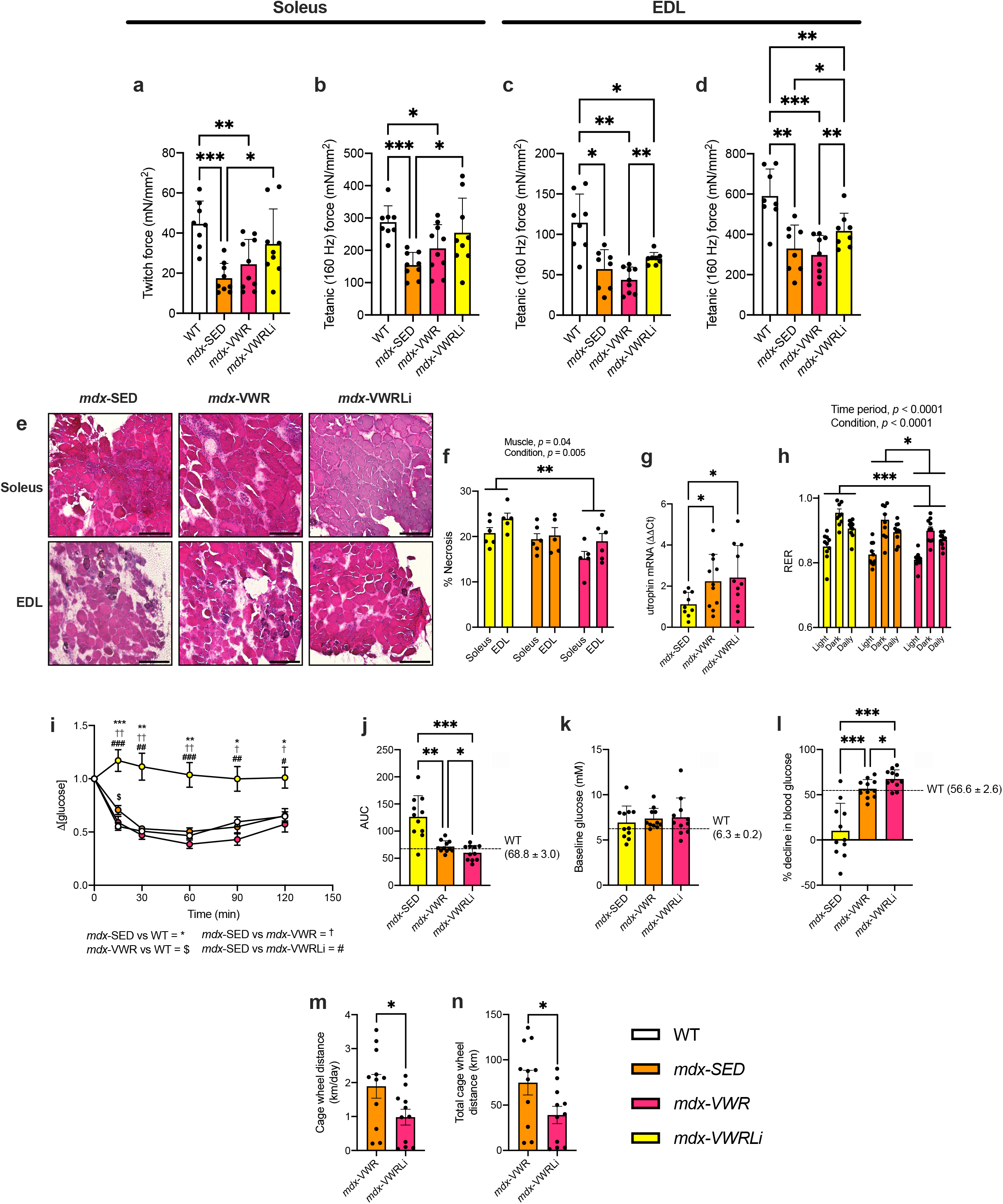
Lithium treatment improves muscle function and insulin sensitivity in *mdx* mice subjected to voluntary wheel running. **a-d)** Twitch and tetanic specific force production in isolated soleus and EDL muscles is improved in the *mdx*-VWRLi group (n = 8-10 per group). **e-f**) Muscle histological analysis with H&E staining shows a significant reduction in % necrosis in soleus and EDL muscles obtained from *mdx*-VWRLi mice vs. *mdx*-SED mice. A significant main effect of muscle type also indicates that EDL muscles have greater % necrosis vs. soleus muscles (n = 5-6 per group). **g)** Utrophin expression is higher in in quadricep muscles from *mdx*-VWR and *mdx-*VWRLi groups when compared with *mdx*-SED mice (n = 9-11 per group). **h)** Respiratory exchange ratio (RER) is lowest in the *mdx*-VWRLi mice when compared to all other *mdx* groups (n = 10 per group). **i-l)** Insulin tolerance tests and corresponding area-under-the-curve (AUC) analysis, baseline glucose levels and % decline in blood glucose in 10-12 week old WT, *mdx*-SED, *mdx*-VWR, and *mdx*-VWRLi mice (n = 11 per group). Glucose traces are displayed relative to baseline glucose levels. AUC were obtained from the normalized traces. There were no differences in baseline glucose levels. **m-n)** Average daily and total distance travelled with a cagewheel was lower in *mdx*-VWRLi mice vs. *mdx*-VWR mice (n = 11 per group). For **a-d** and **g**, a Welch’s one-way ANOVA with a Dunnett’s T3 post-hoc test was used. For **f** and **h**, a two-way ANOVA with a Tukey’s post-hoc test was used; and significant main effects and interactions denoted in the text above. For **i**, a two-way repeated ANOVA was used with a Tukey’s post-hoc test with significance levels denoted in the text below the graph. For **j-l,** a one-way ANOVA with a Tukey’s post-hoc test was used. For **m-n,** a Student’s t-test was used. All values are mean ± SEM. \**p*<0.05, \*\**p*<0.01, and \*\*\**p*<0.001.

With respect to the metabolic phenotype, and similar to that observed with tideglusib treatment (Fig. 2), the *mdx*-VWRLi mice had the lowest RER compared with the *mdx-*SED mice, which is indicative of enhanced lipid oxidation (Fig. 3h). The *mdx-*VWRLi mice were also more insulin sensitive when compared with the other *mdx* groups (Figure 3i-l). Specifically, insulin tolerance tests showed that *mdx*-SED mice were insulin resistant compared with WT, *mdx*-VWR, and *mdx*-VWRLi mice (Fig. 3i). While VWR certainly improved insulin tolerance in *mdx* mice, Li supplementation provided an additive effect, whereby *mdx-*VWRLi mice had the lowest AUC and largest percent decline (∼67%) in blood glucose levels compared with all *mdx* groups and without any differences in baseline glucose levels (Figure 3j-l). Furthermore, the metabolic and contractile benefits of Li were found with half the exercise volume compared with *mdx*-VWR mice (Figure 3m-n). Though we cannot provide an explanation for the lowered exercise volume with Li supplementation, these results show that despite running less, Li supplementation combined with VWR can significantly improve specific force production, lower muscle necrosis, and enhance oxidative lipid metabolism and insulin sensitivity. As some patients with DMD may not be able to engage in adequate amounts of physical activity, our results may be clinically impactful, as they suggest that exercise and treatment with a GSK3 inhibitor may provide synergistic benefits to muscle and metabolic health.

In summary, our data shows that treating the severe DBA/2J mouse model with GSK3 inhibitors either alone or with exercise can improve skeletal muscle health and function, while also improving insulin sensitivity and restoring other metabolic abnormalities. Given that tideglusib is currently being tested for other diseases and that Li is already FDA approved, there is a strong potential to repurpose these drugs for DMD, where clinical trials can look towards not only improving muscle health, but also metabolic health.

## Methods

### Study approval

All animal procedures were approved by the Brock University Animal Care and Utilization Committee (file #17-06-03, 20-07-01, and 22-04-01) and were carried out in accordance with the Canadian Council on Animal Care guidelines.

### Mice and Design

DBA/2J (D2)-*mdx* (stock #001801) and D2 WT (stock #000671) were ordered from Jackson Laboratories at 5-6 weeks of age. For the aged tideglusib experiments, *mdx* mice were ordered from Jackson Laboratories at 7-9 weeks of age with treatment starting at 24-26 weeks of age. All mice were housed in Brock University’s Animal Facility, in an environmentally controlled room with a standard 12:12 hour light-dark cycle and allowed access to food and water *ad libitum*.

### Tideglusib treatment

D2 *mdx* mice were treated with tideglusib at a dose of 10mg/kg/day via oral gavage, whereas a separate vehicle group of D2 *mdx* mice were given 26% peg400, 15% Chremaphor EL and water for 5 days per week for 4 weeks.

### Voluntary wheel running and Li supplementation

D2 *mdx* mice were divided into 3 groups: 1) *mdx*-SED, 2) *mdx*-VWR, and 3) *mdx*-VWRLi based on lean mass (g). The VWR groups were provided with a cage wheel for 6 weeks and all mice were housed in the Techniplast DVC80 with GYM500 software to track cage wheel distance travelled 24/7. The Li-fed *mdx* mice were provided lithium chloride (50 mg/kg/day) in their drinking water with 2 water changes per week for the entire duration of the study.

### RER analysis

Mice were housed in the Promethion Metabolic Screening System for a period of 48 hours to track fuel utilization. The cages are connected to a flow regulator and O_2_ and CO_2_ gas analyzers which measure the volume of carbon dioxide produced (VCO_2_) and the volume of oxygen consumed (VO_2_). Respiratory exchange ratio (RER) was calculated as VCO_2_ divided by VO_2_.

### Insulin tolerance test

Insulin tolerance testing was performed as previously described^44^. Briefly, concentrations of blood glucose were determined by sampling blood from the tail vein at 0, 15, 30, 60, and 120 min following the IP insulin injection (0.5 U/kg) with the use of a hand-held glucometer (Freestyle Lite; Abbott). Plots of the average changes in plasma glucose over time were made for each group, and the average total area under the curve was calculated. Area under the curve is presented in mmol/l × time, and baseline values are set to X = 0. The % decline in blood glucose was calculated by taking the largest drop in blood glucose throughout the 120 min divided by the baseline glucose levels (timepoint 0).

### Hangwire impulse

Whole-body muscle performance was measured using a hangwire test according to the Neuromuscular Disease Network Standard Operating Procedure: DMD_M.2.1.004. All mice were gently placed on the wire situated 12 inches high and were left suspended on the wire until they reached exhaustion and dropped from the wire to the base of the cage. The time they remained suspended was recorded for three trials, separated by a 60s recovery period, and impulse (s*g) was calculated by multiplying the average time suspended by body mass.

### Sample and tissue collection

All mice were euthanized via exsanguination under general anesthetic (vaporized isoflurane) and their tissues and blood were collected. Blood was spun at 5000 x g for 8 minutes (4°C) and serum was collected and stored at −80°C. Extensor digitorum longus (EDL), soleus, plantaris and quadriceps were isolated and stored at −80°C for further analysis.

### Isolated muscle contractility

Soleus and EDL muscles were carefully dissected and mounted onto an Aurora Scientific contractile apparatus (model 305B&701B) to assess muscle force production and fatigue as previously described^14^. Muscles were subjected to a twitch and tetanic (160 Hz) isometric contraction. For data analyses, peak isometric force amplitude (mN) was determined across the range of stimulation frequencies. Peak isometric force was then normalized to muscle cross-sectional area (CSA), which was calculated using the following formula: CSA = m/l*d*(L_f_/L_o_), where m, muscle mass (mg); l, muscle length (mm); d, mammalian skeletal muscle density (1.06 mg/mm^3^)^45^. Lf/Lo is the fiber length-to-muscle length ratio (0.44 for the EDL and 0.71 for the soleus)^46^.

### Serum CK and Li analysis

Serum creatine kinase (CK) activity was measured with a M2 Molecular Device plate reader and a commercially available assay (Cat. #C7522, Pointe Scientific Inc., Canton, MI, USA) fitted onto a 96-well plate and calibrated with a standard curve of purified creatine kinase (Sigma, Oakville, ON, Canada, Cat. 10127566001). Serum Li analysis in the *mdx*-VWRLi mice were measured using the commercially available kit (ab235613, Abcam).

### Immunofluorescent fiber typing

Fiber typing was accomplished through immunofluorescent staining of myosin heavy chain (MHC) isoforms in 10 µm sections of the EDL and soleus muscles as previously described^47^. Slides were imaged using a BioTek Cytation 5 Multimode Plate Reader at 10x magnification using auto-exposure settings for three filters: DAPI, GFP, and Texas Red. Images were then stitched, processed and saved using the Gen5 image processing functions, and then analyzed using ImageJ (NIH) software to estimate fiber type composition and CSA.

### Histological assessment

For EDL and soleus muscles, necrosis was measured using Hematoxylin and Eosin (H&E) staining of EDL muscle samples. All slides were imaged using a BioTek Cytation 5 Multimode Plate Reader. All images were acquired using 10x magnification using auto-exposure settings for the Colour Brightfield setting. Images were then saved and then analyzed using ImageJ (NIH) software to estimate necrotic area. This was done by calculating the total surface area of the muscle, then calculating the sum of the area of all necrosis (areas occupied by collagen and immune cell infiltration) within the sample. This total necrosis was then divided by the total area of the sample.

### mRNA analysis

*mRNA* analyses for utrophin (forward, GGGGAAGATGTGAGAGATTT; reverse, GTGTGGTGAGGAGATACGAT) was accomplished by homogenizing plantaris (*mdx-*vehicle and *mdx*-tideglusib) and quadriceps (*mdx*-SED, *mdx*-VWR, *mdx*-LiVWR) samples with 1mL of TRIzol. A commercially available Qiagen RNeasy kit (Quiagen, Hilden, Germany, ID #74104) and a DNase Max kit (Quiagen, Hilden, Germany, D #15200-50) was then used to isolate RNA. This was then quantified using a NanoVue Plus spectrophotometer (Biochrom Ltd., Cambridge, UK), followed by the use of EcoDry RNA to cDNA (Takara Bio Inc., Kusatsu, Shiga, Japan, ID #639547) reaction tubes and a SimpliAmp Thermal Cylinder (ThermoFisher, MA, USA, ID #A24811) to facilitate the generation of cDNA. Diluted cDNA was then analyzed using a qPCR reaction 96-well plate and the StepOnePlus Real-Time PCR System (ThermoFisher, MA, USA, ID #4376600). Threshold cycle (Ct) values were recorded, and data were analyzed using the ΔΔCt method and reported as a fold-change (2ΔCt), using expression of the Gapdh housekeeping gene as a reference (forward, CGGTGCTGAGTATGTCGTGGAGTC; reverse, GGGGCTAAGCAGTTGGTGGTG; IDT).

### Western blotting

Western blotting was done to assess protein levels of Pax7 and myogenin in EDL muscles from aged *mdx* mice as previously described^15^. Standard SDS-PAGE was performed with BioRad 7-12% TGX gels (4561086; BioRad, Hercules, CA, USA) and polyvinylidine difluoride (PVDF) membranes were used for all proteins (BioRad). All protein blots were blocked with Everyblot buffer (#12010020, BioRad) for 10 min at room temperature and incubated with primary (Pax7, APAX7; myogenin, Myog, both from development studies hybridoma bank) and corresponding anti-mouse horseradish peroxidase secondary antibody (7076, Cell Signaling) prior to imaging with a BioRad ChemiDoc Imager with Immobilon ECL Ultra Western HRP Substrate (WBKLS0500, Sigma-Aldrich). Images were analyzed using BioRad ImageLab software (BioRad) and were normalized to total protein analyzed on a Ponceau stain.

### Statistical analysis

Statistical comparisons were made using a Student’s t-test, a one-way or two-way ANOVA with a Tukey’s post-hoc or a Welch’s ANOVA with a Dunnett’s T3 post-hoc test where appropriate. All statistical analysis were done using Graphpad Prism 8 software. Statistical significance was set to a *p* ≤ 0.05. All values are presented as means ± SEM.

## Data availability

All data reported in this paper will be shared by the corresponding author upon request. This paper does not report original code.

---

**Extended Data Fig. 1. DBA/2J *mdx* mice (10-12 weeks) have elevated respiratory exchange ratio (RER) when compared with healthy WT mice.** A two-way ANOVA was used to examine the main effects of time period and genotype. Significant main effects are denoted in the text above.

## Acknowledgements

This work was supported by an unrestricted Brock University Grant, a Canada Research Chair (Tier 2) award to VAF, and by a Brock-Niagara Validation Prototyping and Manufacturing Applied Project Grant in partnership with AMO-Pharma. BMM was supported by an NSERC USRA. SIH was supported by a NSERC Doctoral Vanier Award, JLB was supported by a CIHR CGS-D award, and BH was supported by an Ontario Graduate Scholarship. We thank Tonya Coulthard (Scintica Instruments) for her help with high frequency ultrasound analysis. We thank the animal care staff at Brock University for their assistance with animal care and handling.

## Author contributions

BMM - Concept design, writing and drafting the manuscript, figure creation, data collection, data interpretation, particularly on skeletal muscle function.

BLH – Data collection, data interpretation, reviewing and revising the manuscript

JLB – Data collection, data interpretation, reviewing and revising the manuscript

KCW – Data collection, data interpretation, reviewing and revising the manuscript

MSG – Data collection, data interpretation, reviewing and revising the manuscript

RWB – Data collection, data interpretation, reviewing and revising the manuscript

CJFW – Data collection, data interpretation, reviewing and revising the manuscript

SS – Data collection, data interpretation, reviewing and revising the manuscript

SIH – Data collection, data interpretation, reviewing and revising the manuscript

LJW – Data collection, data interpretation, reviewing and revising the manuscript

RWE – Data collection, data interpretation, reviewing and revising the manuscript

AM – Data collection, data interpretation, reviewing and revising the manuscript

RV – Provision of reagents and resources, reviewing and revising the manuscript

BDR – Provision of reagents and resources, reviewing and revising the manuscript

AJM – Provision of reagents and resources, reviewing and revising the manuscript

REKM – Provision of reagents and resources, reviewing and revising the manuscript

VAF – Concept design, supervision, provision of reagents and resources, acquiring funding, writing and drafting the manuscript

## Competing interests

A portion of this project, specifically the aged (28-30 weeks) tideglusib experiments, were funded by AMO-Pharma in partnership with a Brock-Niagara Validation Prototyping and Manufacturing Applied Project Grant (1:2 monetary ratio). AMO-Pharma also provided the tideglusib for the aged tideglusib experiments.

## Notes

### Summary of Updates

This version focuses solely on skeletal muscle data.

