## Supplementary figures and images for "GSK3 inhibition improves skeletal muscle function and whole-body metabolism in the severe DBA/2J *mdx* mouse model"

### Extended Data Fig. 1

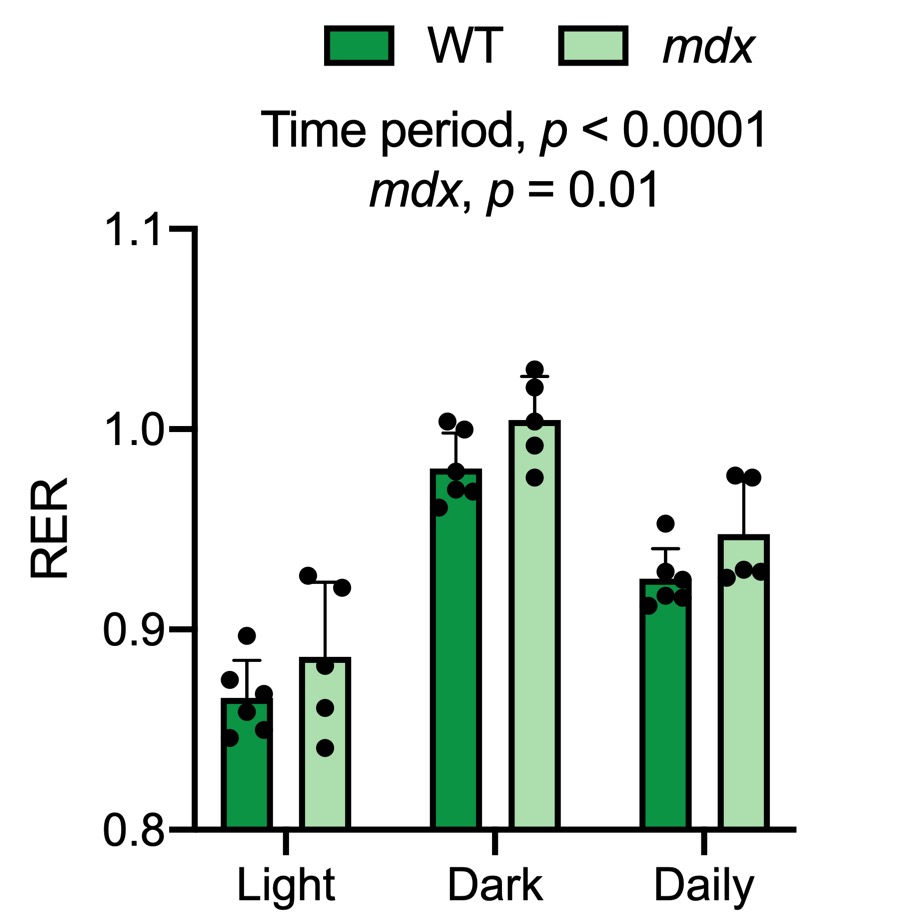
